# PatternExtract: A facile, scalable pipeline for point pattern generation from spatial imaging data

**DOI:** 10.1101/2025.06.25.661424

**Authors:** Shruti Sridhar, Gayatri Kumar, Siddham Jasoria, Ziwei Meng, Patrick Jaynes, Vaibhav Rajan, David Scott, Claudio Tripodo, Kasthuri Kannan, Anand D Jeyesekharan

## Abstract

We present PatternExtract, an open-source pipeline that generates accurate spatial point patterns from RGB pathology images and cell coordinate data without relying on composite channels or proprietary software. Using a novel two-kernel tissue segmentation method combined with automated pixel classification in QuPath, PatternExtract precisely excludes tissue artifacts such as necrosis and blood vessels to define spatial windows. Optimized on 568 diffuse large B-cell lymphoma images and validated on an independent cohort, the pipeline enables robust spatial analyses.

## Introduction

Spatial biology is increasingly integral to cancer research. With the advent of methods to provide molecular information at single cell resolution, preserving spatial detail in sections of cancer tissue, it is now possible to evaluate inter-relationships between malignant cells and the tumor microenvironment (TME). However, most published spatial analyses in cancer rely on nearest neighbor distance measures, which provide a simplified, global summary of cell proximities. However, such aggregated measures may obscure spatial heterogeneity in spatial distribution of cells within the region of interest (ROI) and limit biological insight.

A more robust approach is through spatial point pattern analysis. This mathematical framework is a method to study the relationships between data points in 2-dimensional space, and is well-established in fields such as ecology (Baddeley et al., 2015; Illian et al., 2008), where for instance it is used to examine interactions between plant species. It is also widely used in geographic information systems (GIS) to mathematically quantify underlying relationships between spatial features. Representing data as two- or three-dimensional spatial point patterns enables the application of rigorous mathematical functions in various spatio-temporal studies. These methods can quantitatively distinguish patterns such as clustering from complete spatial randomness (CSR) (Wiegand & Moloney, 2014). Mathematically, CSR corresponds to a Poisson distribution and helps to infer whether spatial features are randomly distributed, ordered, or clustered.Drawing from the utility of spatial point pattern analysis in ecology, these approaches are potentially valuable to understand the ecosystem of cancer using multiplexed immunohistochemistry (IHC) or multiplexed immunofluorescence (mIF) data, which provide spatial coordinates for biomarkers at single-cell resolution. The geospatial R package **spatstat** (Baddeley et al., 2015) offers a comprehensive set of tools to fit such models and simulate spatial point processes, and mathematical models like Gibbs processes allow multi-scale modeling of cellular inhibition and regularity within the tumor microenvironment (TME), such as homotypic cell inhibition (Kumar et al., 2023). However, reliable inference from these models requires well-defined spatial windows that exclude non-cellular regions such as necrosis or tissue artifacts. Existing spatstat boundary functions typically approximate the ROI using convex hulls or rectangular windows, which can include artifacts such as blood vessels or tissue damage, posing a challenge for accurate spatial analyses.

To address these challenges, we developed a robust pipeline *PatternExtract* to generate precise spatial point patterns directly from RGB pathology images, without requiring composite marker channel images or proprietary software. The pipeline uses spatial coordinates to create a two-kernel mask per image, automatically excluding regions of tissue loss and blood vessels, a critical step for analyzing cancer tissue sections, particularly when in the tissue microarray (TMA) format. This pipeline leverages QuPath, an open-source digital pathology platform, to display images and annotations for user review, while automating pixel classification to adaptively exclude artifacts.

We developed and optimized this pipeline on a large cohort of 568 images from 274 patients of diffuse large B-cell lymphoma (DLBCL). The pipeline consistently performs well, generating accurate point patterns independent of marker type. To validate its generalizability and explore biological applications, we applied the pipeline to an independent set of 20 images (n = 20 patients) from another cohort, demonstrating its utility for spatial analysis across diverse datasets and enabling reliable inference in TME studies of DLBCL.

## Methods

### Samples and Dataset

Patients diagnosed with DLBCL (*n* = 274), tissue microarray (TMA) format were obtained from British Columbia Cancer Agency (*Ennishi et al)*. Additional DLBCL TMA were obtained from the NUH [approved by the Singapore NHG Domain Specific Review Board B study protocol (2015/00176)]. Samples were obtained through IRB-approved ethics protocols, with written informed consent from the patients, or with IRB-approved waivers of consent where applicable in accordance with the ethical guidelines of the Declaration of Helsinki. Material transfer agreements from all providing institutions were incorporated into the framework of an NUS IRB-approved translational study (H-19-055E).

### Immunofluorescence staining

Quantitative mfIHC was performed using sequential OPAL-TSA staining. Images were acquired using the Vectra 2 imager and analyzed using QuPath. DAPI nuclear staining and CD20 membrane staining were used to segment cells. The mean membrane intensity per cell was captured for CD20; and the mean nuclear intensity per cell for Ki67. For each image, cells with a marker intensity above a given intensity threshold for that image were declared to be positive for that marker. A pathologist manually inspected each image to determine a reliable threshold for each marker that resulted in minimal false-positive and false-negative assignments. The cohorts were evaluated in a tissue microarray (TMA) format; depending on tissue availability and between 1 and 9 high-power 700 × 500 μm imaging fields were captured and evaluated per patient. Following this the spatial pipeline was applied to derive spatial point patterns.

### Generating tissue segmentation masks

Spatial windows can be generated for any image that has an accurate segmentation mask. The proprietary software InForm generates a tissue segmentation mask that can be used for this purpose. To build such a mask using the RGB images and spatial coordinates we used an image processing package called *OpenCV*. The pipeline can be generalized to all images if the coordinates with marker information is provided as a *csv* file. Our method overlays points on the RGB images factoring in how the coordinates scale with reference to the image. Using a two kernel approach (Fig 1A) with differing intensity and radius we were able to obtain an accurate mask for 568 RGB images without the software generated tissue segmentation mask. This tissue mask can be used to train a pixel classifier in QuPath *(Bankhead et al*., *2017)*, an open source digital pathology tool.

**Figure 1.**
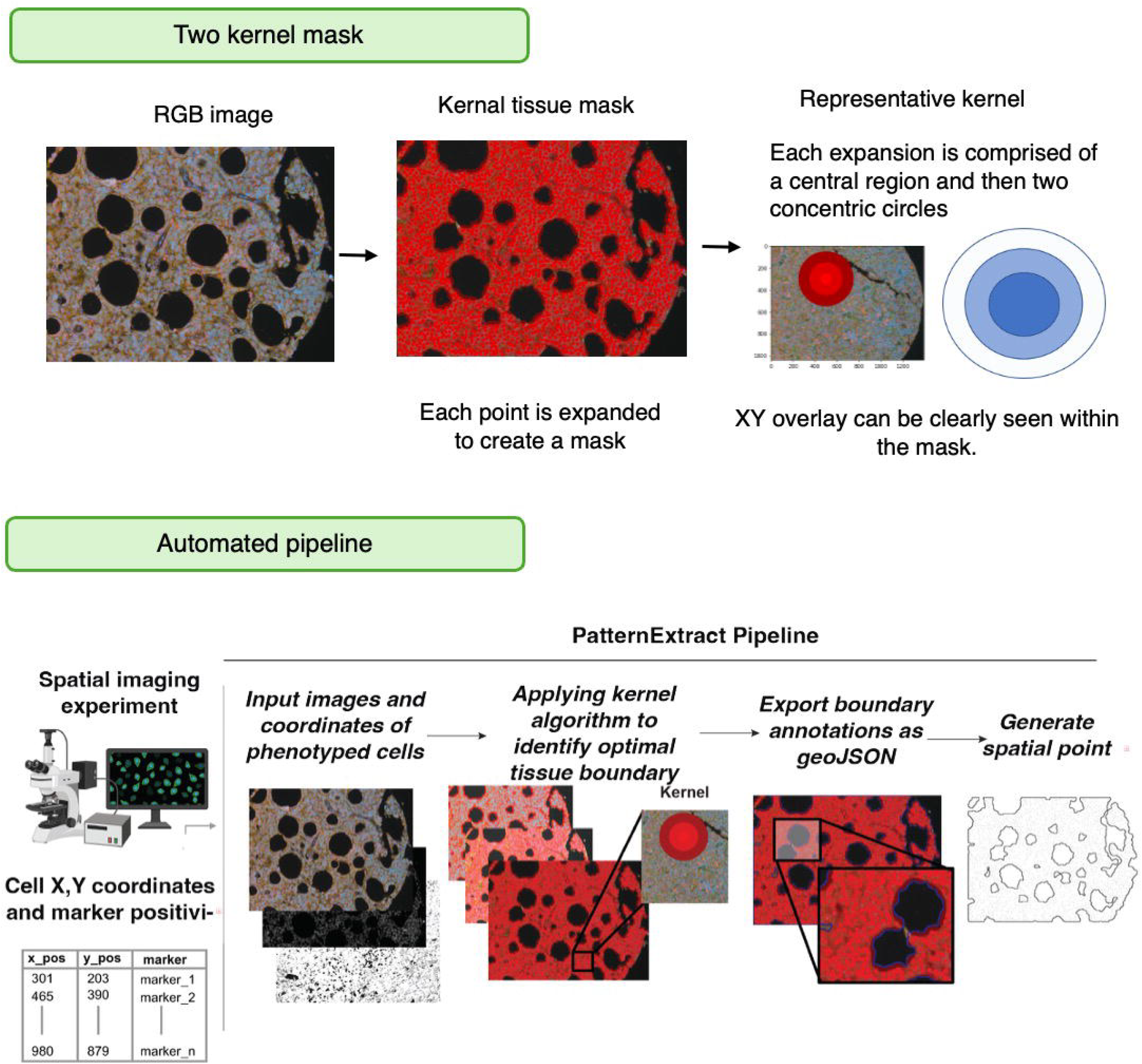
Overview of the PatternExtract pipeline for generating spatial point patterns from multiplex imaging data. (Top) A two-kernel masking approach is used to define tissue regions from RGB images. Each detected cell (via XY coordinates) is expanded into two concentric circles to create a composite tissue mask that highlights true tissue regions while excluding voids. (Bottom) The automated PatternExtract pipeline starts with spatial imaging data and cell coordinate tables. Input RGB images and corresponding cell coordinates are processed using the two-kernel algorithm to create accurate tissue boundaries. These boundaries are exported as GeoJSON annotations from QuPath, replacing default convex hulls. Final spatial point patterns are generated within these refined boundaries for downstream spatial analysis.

### Pixel classifier in QuPath for generating annotation: fine tuning parameters

The QuPath tool includes a built-in pixel classifier that can be trained on images to generate tissue segmentation masks. The classifier uses the *create thresholder* function, along with tunable parameters such as *minimum object size, minimum hole size*, a residual channel with a *Gaussian prefilter*, and the *Full resolution* setting *(downsample = 1*) to produce GeoJSON annotations. During training, the threshold categories *“Unclassified”* and *“Region*^*\**^*”* are used to annotate the entire image. These GeoJSON annotations contain a dictionary list of XY coordinates, which can then be imported to define spatial windows that accurately capture the tissue boundaries and accommodate heterogeneity across tissue sections.

After training, the classifier is first tested on a single image. Based on the optimized parameters, a Groovy script is generated to apply the classifier across the entire project and to automate the export of the GeoJSON annotations. The resulting annotated images are manually reviewed to verify that excluded regions are correctly identified and not misclassified. When necessary, the *“Fill Holes”* function can be applied to correct any erroneous exclusions in the annotations.

### Importing annotations for spatial point pattern windows

The *csv* file is used to generate marked planar point patterns with *marks* from the *spatstat* toolbox. *Marks* are the combination of biomarker expressions for each cell in the TMA image. Spatial point patterns are generated for each image and then taken forward for spatial analysis. The spatial window we used has an inbuilt function *convexhull* which provides an approximation based boundary around the cells. The exported geojson annotations for each image are plugged in to replace the convex window. The percentage of points that lie outside the boundary and excluded by the point pattern is used as a measure of the accuracy of the pipeline. This step involved iterations to fix regions where cells were on the edge, outside the boundary. The points excluded due to *minimum object size* set to 5000 px^2^ were identified and the threshold was relaxed.

### Automating the pipeline for the project

The different steps detailed above used to generate spatial point patterns from any type of TMA using open source tools have been consolidated into a pipeline. A single line command can be used to execute the python file which requires three system arguments which are the paths to the csv files, the raw RGB files and the folder to write the exported masks. The python file calls the other scripts for generating the point patterns. Also, it runs a QuPath executable which pops up the GUI viewer to review the annotations generated by the groovy script with the parameters used on the benchmarking dataset (Fig 1B).

The complete codebase for the pipeline is available on GitHub at: https://github.com/shrutisridhar99/PatternExtract

## Results

### Two kernel tissue segmentation mask for TMA images

We generated tissue segmentation masks with a two kernel approach by overlaying the spatial coordinates on the 568 image cohort from BCA. This cohort consisted of RGB images of multi fluorescence immunohistochemistry without an available inForm tissue segmentation mask. Only the red,green,blue channels are available for the RGB images when viewed in QuPath as brightfield fluorescence. The spatial coordinates of the single cell expression of different oncogenes is used to develop an ad hoc tissue mask. Generating the spatial point patterns relies on the contours of the tissues and excluding the tissue artifacts such as a folded or damaged tissue. We developed a pipeline for spatial analysis of this cohort by developing the tissue mask using the spatial coordinates and an image processing package, namely OpenCV. Overlaying the coordinates was a starting point and after several iterations of point size and radii we arrived at the two kernel approach to define each cell as two concentric circles (Fig 2). The larger circle with lower red channel intensity circumscribes the smaller circle with higher intensity (Fig 1A). This approach defined the tissue contours and the single cell positions without dependence on proprietary software. The tissue mask generated using this method was used to train the pixel classifier in QuPath.

**Figure 2.**
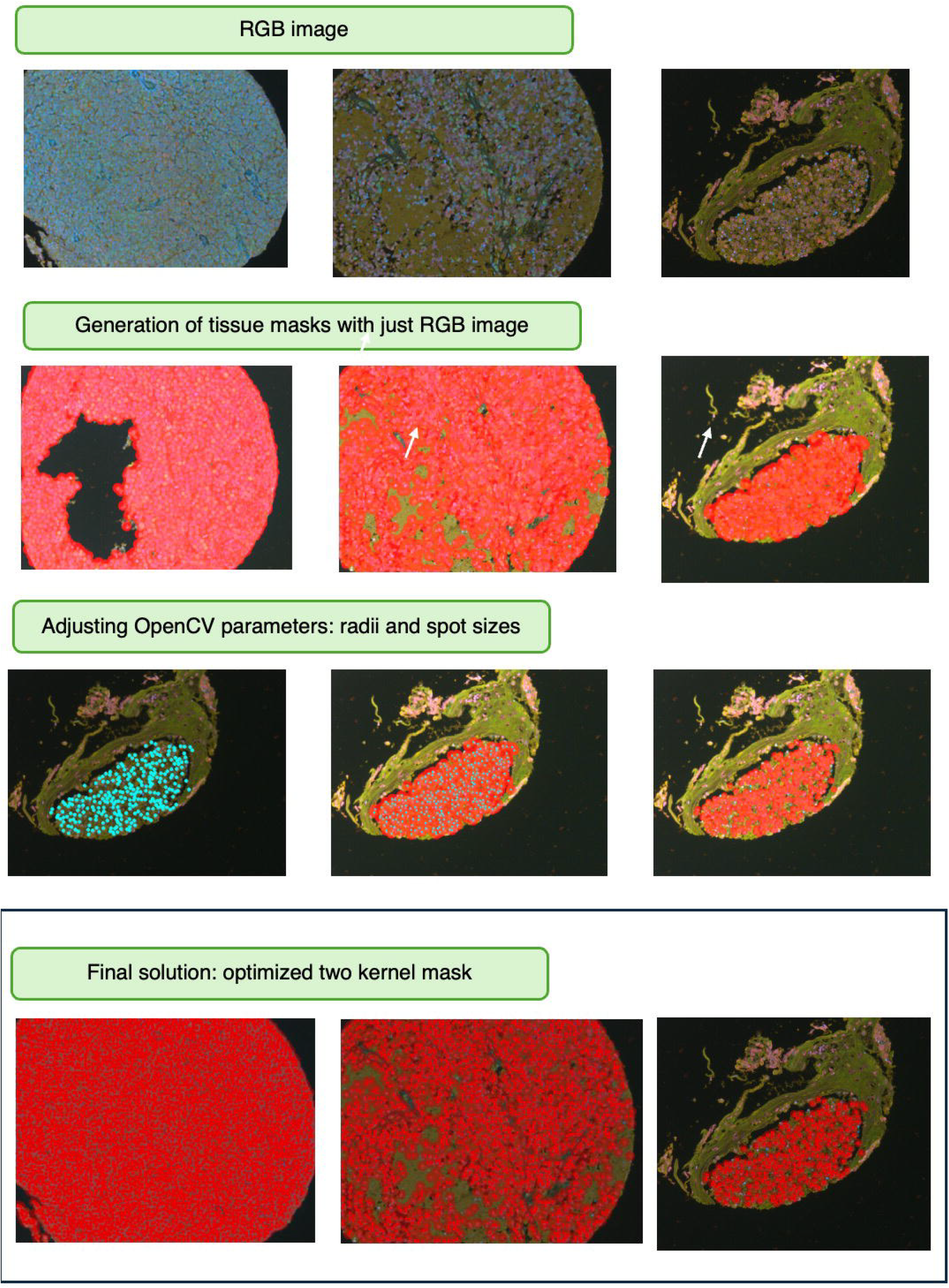
Figure 2: Development and optimization of the two-kernel tissue masking approach from RGB images. (Top row) Representative RGB images from multiplex immunofluorescence-stained tissue sections used as input for mask generation. (Second row) Initial tissue masks generated directly from RGB images using OpenCV, highlighting issues such as tissue loss (white arrows) and inclusion of non-tissue regions. (Third row) Parameter tuning in OpenCV, specifically radii and spot sizes, improves the accuracy of point overlay and mask coverage across heterogeneous tissue regions. (Bottom row) Final optimized two-kernel mask output, where the concentric circles accurately delineate tissue boundaries and exclude artifacts, enabling precise spatial point pattern generation without requiring proprietary software.

### Tissue artifacts: tissue damage and innervating blood vessels

One of the challenges we face with large cohorts is the inherent tissue heterogeneity such as clumps of cells interspersed with regions without cells. Occasionally, the sections have tissue loss with multiple holes in the core. It is also imperative to exclude the areas of the image without cells as they could be necrotic regions. In some cases the sections show intact tissue with no folding, however they have innervating blood vessels leading to circular regions in the tissue without cells. The reddish orange color of the RGB images without cells is also identified by the pixel classifier in spite of the two kernel masks generated by our method. These features in the data pose a challenge in determining the parameters for the classifier such as *minimum object size*. This is because a large object size can lead to the exclusion of clumps of cells in regions less than 5000 px^2^. Relaxing this parameter will include artifacts because of the red-orange background. We developed a code frame that can be generalized across tissue regions accounting for heterogeneity, by experimenting with parameters for the classifier and by converting the RGB images to grayscale to avoid false positives and setting an optimal minimum hole size (Fig 3A). These are shown in the examples below where regions with fibrotic tissue and holes where the final point pattern accurately avoids these areas (Fig 3B).

**Figure 3.**
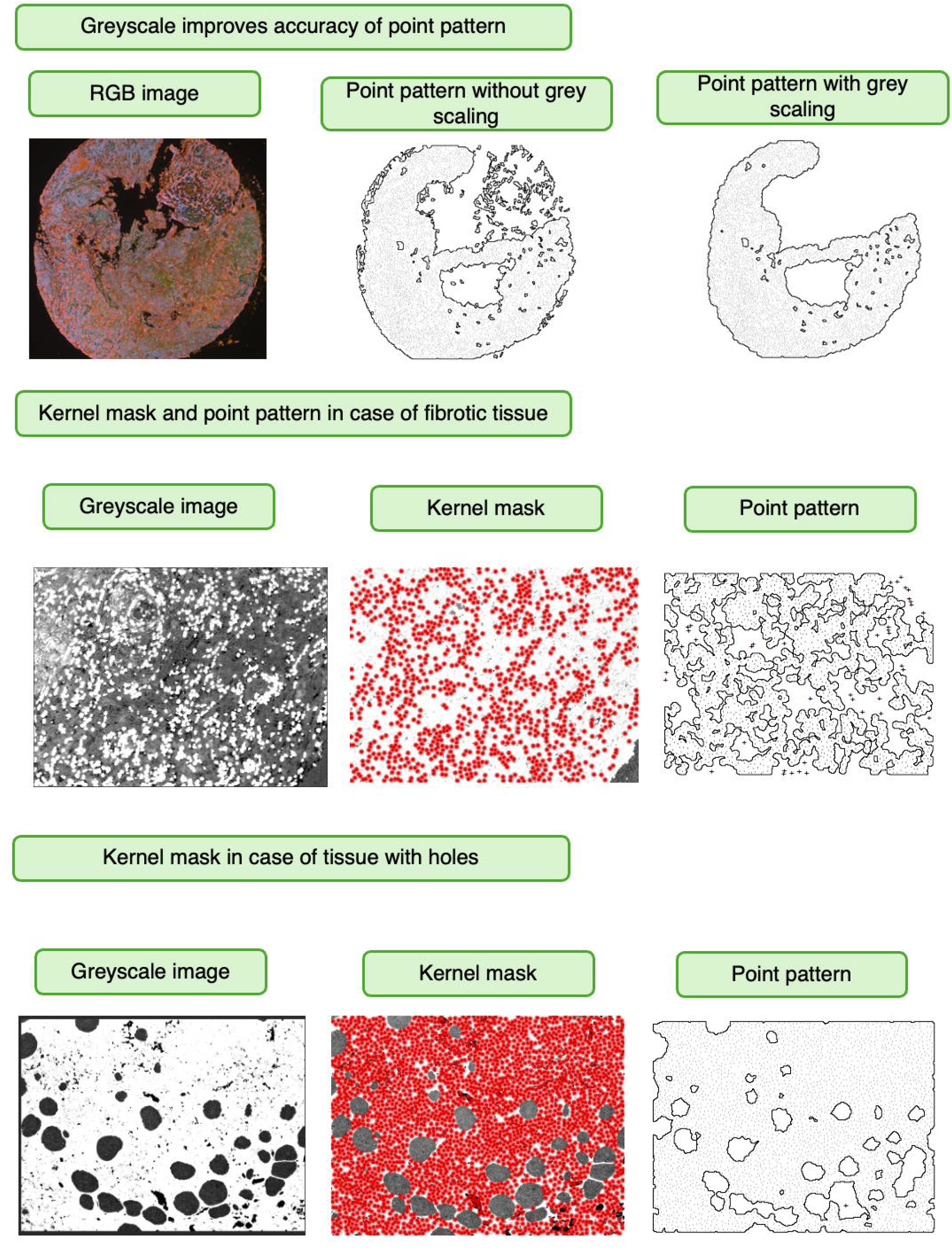
Refinement of point pattern accuracy using grayscale transformation and kernel mask adjustments in challenging tissue contexts. (Top row) Point pattern generation is improved by converting RGB images to grayscale. Without grayscale conversion, artifacts and non-tissue regions are mistakenly included (middle). Grayscale preprocessing (right) enhances boundary accuracy and spatial pattern fidelity. (Middle row) In fibrotic tissue, dense and heterogeneous staining complicates segmentation. Grayscale transformation (left) facilitates cleaner kernel-based masking (middle), producing a more accurate spatial point pattern (right). (Bottom row) For tissues with holes or structural voids, grayscale images (left) allow the kernel mask (middle) to effectively exclude empty regions, resulting in a cleaner and biologically accurate point pattern (right). These examples highlight the adaptability of the PatternExtract pipeline in processing varied tissue morphologies and improving spatial analysis through image preprocessing and dynamic kernel tuning.

We incorporated the parameters in a groovy script for the entire project to generate annotations and export them as individual *geojson* files. The geojson annotations are imported in R script and replace the *convexhull* spatial window of the point patterns with the QuPath exported annotations.

The Qupath software includes an in-built command line script editor which we used to process the set of images (568) with the 2 kernel overlaid spatial coordinates and automate this step of the process. The QuPath pixel classifier is able to identify the tissue boundaries and also the tissue regions devoid of cells and exclude void regions without tissue in the TMA. These are then plugged into an R script that builds the spatial point patterns with marks for each of the images. The default convex window is replaced by the geojson annotations and accurate spatial point pattern windows are obtained for the cohort without the assistance of any proprietary software.

### Performance evaluation of spatial pipeline

The performance of the pipeline can be visually assessed by the accuracy of the QuPath annotations in the viewer. However, a quantitative measure of performance was tabulated by calculating the percentage of points lying outside the spatial boundary imported from the QuPath annotations. This is a reflection of the quality of the tissue mask and thereby the accuracy of annotations. In some cases the annotations imported into the R script for replacing the default spatial point pattern window can have a scaling issue such that a shift in the mask is seen for a few images. This was addressed by replacing the scaling factor applied to adjust the default window xy min-max limits relative to the *xy* limits of the new spatial window. This can be seen in Fig. 4A and the improvement with the new scaling can be seen on the right Fig. 4B. The number of images with points excluded also decreased with the new scaling Fig 4C.

**Figure 4.**
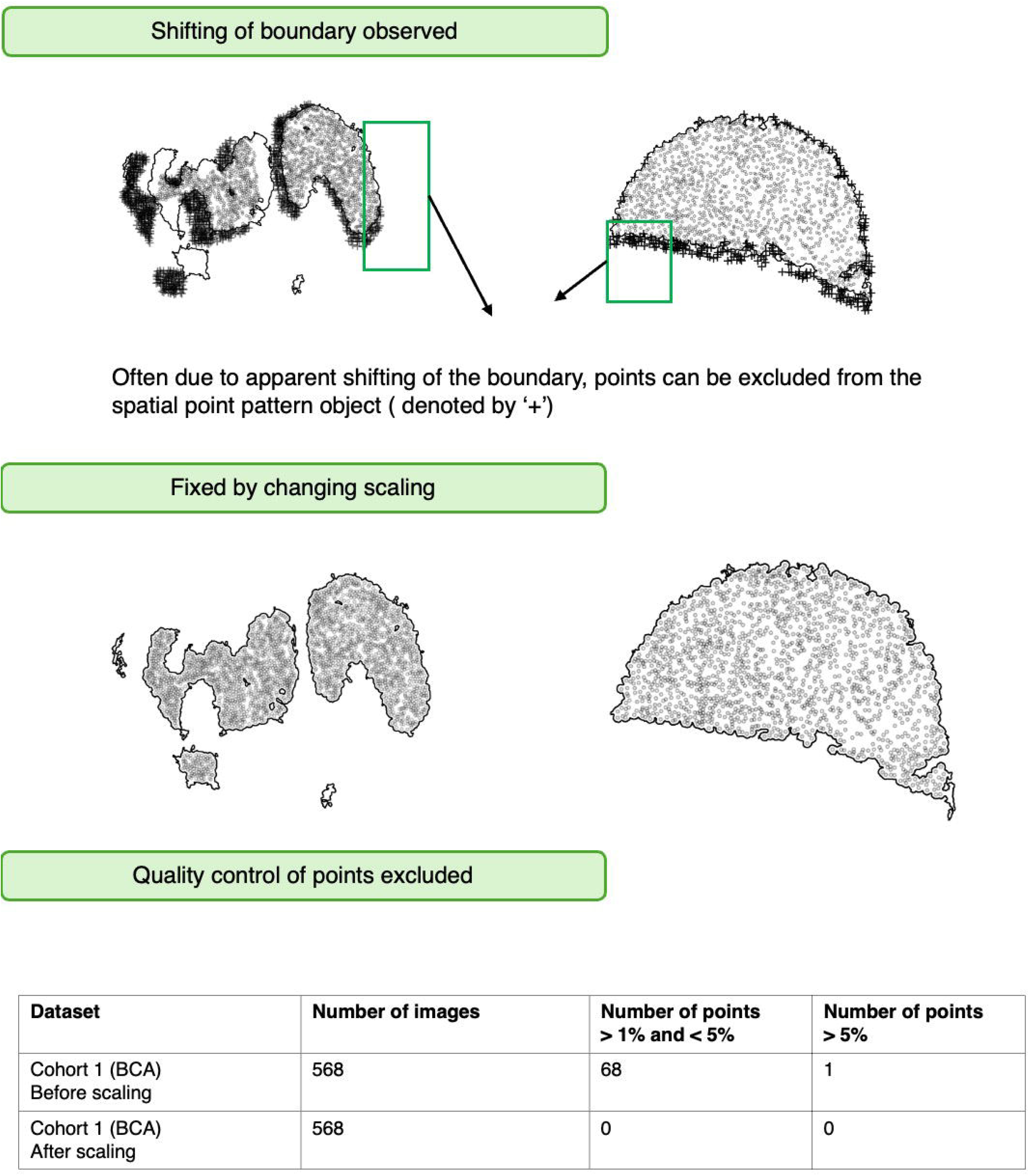
Quality control pipeline for spatial point pattern extraction addressing boundary shifting artifacts and ensuring data integrity. (Top row) Boundary shifting artifacts can cause exclusion of legitimate points from the spatial point pattern object. The highlighted regions show areas where apparent boundary movement leads to point exclusion, compromising the completeness of the extracted pattern. (Middle row) The scaling adjustment approach resolves boundary shifting issues by optimizing the spatial scale parameters. This step ensures consistent boundary detection across the tissue section, eliminating artificial point exclusions while preserving the underlying spatial structure. (Bottom row) The quality control summary table demonstrates the effectiveness of the scaling correction approach. Before scaling adjustment, Cohort 1 (BCA) showed 68 points with moderate exclusion rates (>1% and <5%) and 1 point with high exclusion (>5%) across 568 images. After implementing the scaling correction, all exclusion artifacts were eliminated (0 points in both categories), confirming the robustness of the quality control pipeline.

### Spatial pipeline standalone package

The workflow used for generating spatial point patterns consists of the following parts – The python script with using the OpenCV package to manipulate the kernel size and intensity of the spatial coordinates overlaid on the images. A QuPath groovy script to automate the import and export of the images into the project folder for QuPath and the parameter thresholds for the pixel classifier. An R script to build the spatial point patterns from coordinates and save different oncogene co-expression as marks within a default convex window. Replacing the convex window with the accurate geojson annotations exported from QuPath and comparing the two. Obtaining measures of improvement in terms of minimizing the number of points excluded while generating accurate spatial point pattern windows. Using an R script to calculate the pairwise clustering and its deviation from complete spatial randomness (i.e. a Poisson distribution) using the *pcf* function. For the scientific community consisting of cancer biologists, pathologists and clinicians with limited domain knowledge in programming, the individual steps of the above mentioned spatial analysis scripts have been consolidated into a standalone package that can be executed in a facile manner. The package supports the visualization of the images with the annotations through a pop-up QuPath window. The user can tweak and refine the annotations built by the classifier in QuPath at this intermediate step. The pipeline can be run on any image with a corresponding *csv* file containing the spatial coordinates.

### Biological application of point patterns

We investigated the spatial distribution of Ki67-positive cells in Diffuse Large B-cell Lymphoma (DLBCL) (n = 20) to illustrate the downstream utility of spatial point pattern analysis Fig 5A. Ki67, a well-established marker of proliferation *(Miller et al*., *2017; Keane et al*., *2015)*, has been previously studied as a prognostic factor in DLBCL. To assess its spatial organization, we employed point pattern analysis with a focus on deviations from complete spatial randomness (CSR), using the pair correlation function (PCF).

**Figure 5.**
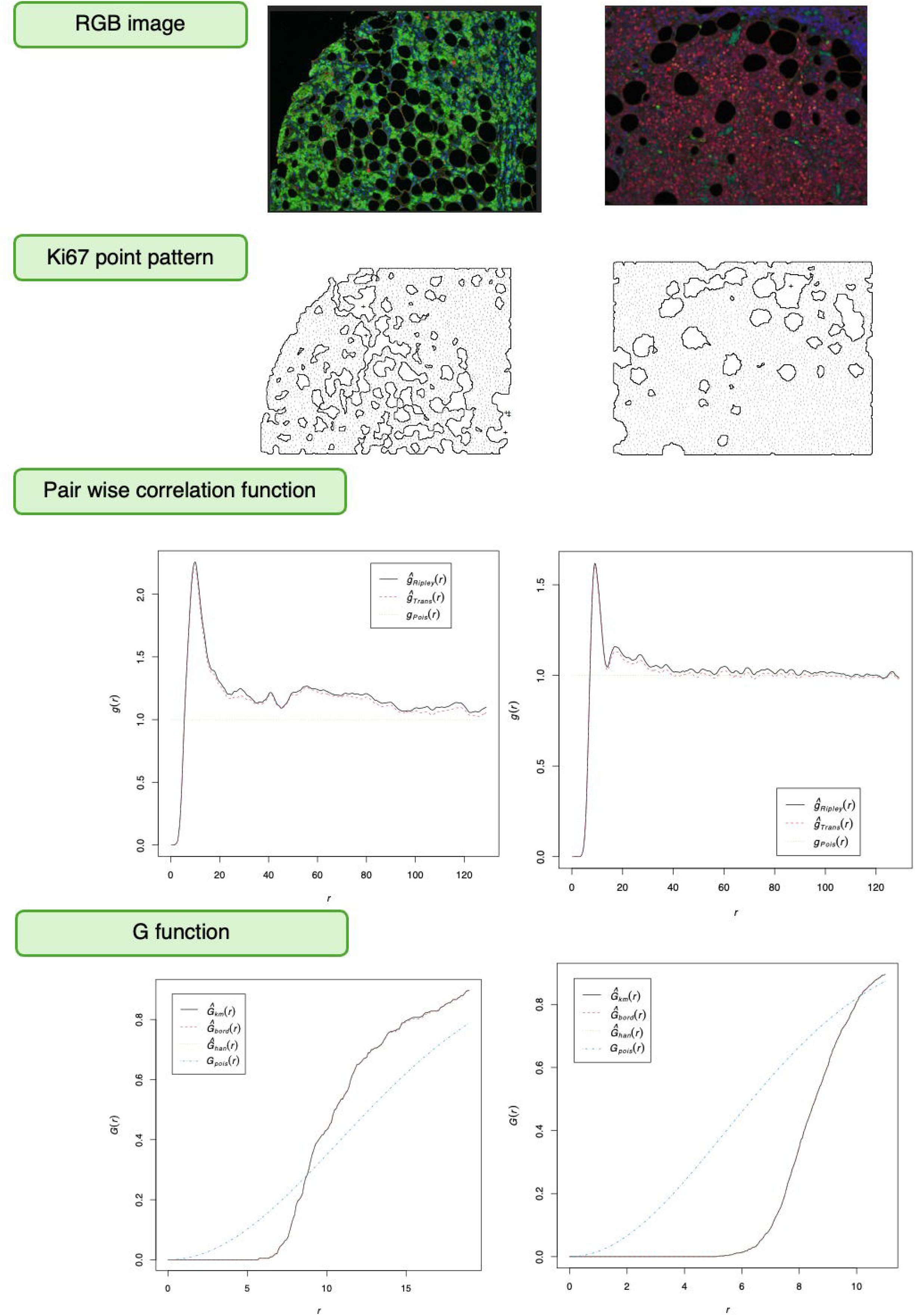
Spatial analysis workflow demonstrating the progression from tissue imaging to quantitative spatial statistics for point pattern characterization. (Top row) RGB fluorescence microscopy images showing Ki67 staining of representative images. (Second row) Ki67 point pattern extraction using PatternExtract. (Third row) Pair-wise correlation function analysis (g(r)) quantifying spatial clustering and dispersion patterns at different length scales. The left panel shows strong short-range correlation, indicating clustered spatial organization. The right panel demonstrates more uniform spatial distribution with sustained correlation values, reflecting different underlying tissue organization patterns. (Bottom row) G function cumulative distribution analysis measuring nearest-neighbor distances to characterize spatial point processes. The curves represent different statistical models (Ĝobs(r), Ĝenv(r), Ĝpois(r), Ĝcsr(r)) comparing observed patterns against theoretical distributions, enabling classification of spatial arrangements as clustered, dispersed, or random relative to complete spatial randomness expectations.

The PCF was used to quantify local clustering by measuring the probability of finding a pair of cells at a given distance compared to a random distribution. Our analysis revealed a marked increase in clustering of CD20□ Ki67□ cells at short distances, indicating localized proliferation Fig 5B.

To further characterize the spatial arrangement of proliferating cells, we analyzed the G function, which measures the cumulative distribution of nearest-neighbor distances. Unlike the PCF results that suggested local clustering, the G function revealed a deviation below the theoretical expectation under spatial randomness, particularly at short intercellular distances Fig 5C. This suggests a degree of local inhibition or spacing among CD20□Ki67□ cells. Together, these findings highlight the complexity of spatial interactions, emphasizing the importance of using multiple complementary spatial statistics to fully capture both aggregation and exclusion patterns. This example highlights the value of spatial point pattern analysis in revealing microarchitectural features of the tumor microenvironment and demonstrates the applicability of our analytical pipeline.

## Discussion

We introduce *PatternExtract*, a robust, open-source pipeline to generate biologically accurate spatial point patterns directly from RGB images and coordinate data. A key strength of *PatternExtract* lies in its ability to automate tissue mask generation using a two-kernel approach that captures tissue geometry and density without relying on fluorescence composite channels or vendor-specific image formats. This feature is particularly advantageous in retrospective or collaborative studies where only RGB images and cell coordinate files are available. Moreover, by exporting high-fidelity GeoJSON annotations from QuPath the pipeline mitigates the inclusion of non-tissue artifacts and voids, improving the accuracy of spatial summary functions such as the pair correlation function (PCF) and G-function. *PatternExtract* is also suitable for integration into larger image analysis pipelines, including those incorporating spatial transcriptomics or multimodal imaging. Future iterations could also extend support for batch inference of spatial interactions across multiple marker pairs and incorporate deep learning-based segmentation to further improve robustness. Overall, *PatternExtract* enhances the reproducibility and biological fidelity of spatial statistics in tumor immunology, offering a valuable tool for spatial analysis at scale.

